# Structure Of Eukaryotic Type III Glutamine Synthetase Lacking Ring-Ring Quaternary Contacts

**DOI:** 10.1101/2022.04.05.486999

**Authors:** Irina V. Novikova, Samantha M. Powell, Chuck R. Smallwood, Samuel O. Purvine, Mowei Zhou, James E. Evans

## Abstract

There are three major classes of the enzyme Glutamine Synthetase (GS), denoted as GSI, GSII and GSIII in both prokaryotes and eukaryotes, that share widely conserved active-site residues but vary in average protein length, oligomeric assembly state and cofactor requirements. Key structures have been determined for all prokaryotic and eukaryotic GS classes except the GSIII class of eukaryotic origin, where even basic questions about its functional oligomeric state are unresolved. Here, we report the 3.2Å cryo-EM structure of eukaryotic GSIII from the ancient organism of green algal lineage *Ostreococcus tauri*. Using a combined structural and biochemical approach, we demonstrate that *O. tauri* GSIII self-assembles and functions exclusively as a single hexameric ring. This is unlike most GSs in other classes, where they form double-stacked ring assemblies instead. Major structural differences in the ring-ring stacking across the classes suggest evolutionary pressure may have favored higher-order interactions that might be beneficial but are not necessarily required for the catalytic GS performance.

## Introduction

In all life forms, glutamine is one of the most abundant amino acids, and plays a vital role in many metabolic processes such as protein biosynthesis, purine biosynthesis, lipid biosynthesis, regulation of homeostasis and transport and storage of nitrogen. The amino acid glutamine is synthesized by a large family of enzymes called Glutamine Synthetases (GSs) that utilize inorganic ammonium and glutamate as substrates. As a result, GSs are critical for organism survival and, therefore, have become favored targets for rational drug design and gene editing to either inhibit or limit the nitrogen production in a desired organism. For example, since nitrogen is a main limiting factor for plant growth, the agricultural industry has widely used phosphinothricin (PPT), an irreversible GS inhibitor^1^, as an herbicide since the 1970s. Similarly, drug screening trials against tuberculosis GS proteins are undergoing^2-4^. Human GS is also being considered as a potential drug target to regulate several neurological disorders^5,6^.

Class I GS (GSI), with molecular weights averaging 45 kDa, are considered primarily prokaryotic enzymes yet close homologs of this class exist in plants and mammals^7-9^. Several crystallographic studies discovered that the GSI enzymes tightly self-organize into dodecamers with D6 symmetry^10,11^. This dodecameric complex is composed of two hexameric rings stacked in a “face-to-face” orientation that interacts quite tightly via formation of 6 ß-sheets built with internal loops and 12 helices extending from the C-terminal of each subunit. Each of the helices is inserted into a hydrophobic cavity of another subunit located on the opposite ring. Thus, every subunit pair (out of 6) is stabilized by 37 hydrophobic and 36 hydrogen bonding interactions^12^.

Class II GSs (GSII) are considered eukaryotic-specific but instances of them are also found in some families of bacteria^13,14^. Individual GSII subunits are slightly smaller in size (around 35-40 kDa), where ten subunits participate in the protein complex formation, forming two stacked pentameric rings with D5 symmetry^12,15,16^ (Fig.S1). On top of the variance in complex stoichiometry between classes I and II, the ring-ring stacking interactions in the quaternary fold differs remarkably between GSI and GSII isoforms^10,12^. In comparison to GSI, GSII complexes are only lightly stabilized with 4 hydrophobic and 2 hydrogen bonding interactions per every subunit pair (out of 5) due to a truncated C-terminus and internal amino acid deletions. Additional evidence, demonstrating the weakness of the ring-ring associations in GSII enzymes, was later obtained through cryo-EM data of the plant GSII from *M. truncatula*^17^. In that study, cryo-EM was able to capture intermediate “swinging” stages of ring-ring quaternary assembly/disassembly, suggesting a rather dynamic and weak nature of these contacts. The latter was additionally supported by other biochemical experiments of the same plant revealing that GSII exists as and is functional as a single pentameric ring in root nodules^18^. An active single-ring GSII complex has been also found in the roots of *Beta vulgarism*^19^.

Despite being widespread amongst bacteria, algae and fungi, the class III of GS (GSIII) is the most sequence diverged from classes I/II and the least characterized to date. GSIII enzymes are much larger with an average mass of 70-80 kDa, and they share less than 10% global sequence identity with other GS classes^20^. GSIII proteins exist both in bacteria and eukaryotes^21,22^. A single crystal structure is currently available for prokaryotic GSIII from *B. fragilis*^23^. The structure showed a two-ring dodecameric assembly similar to GSI, the interfaces responsible for ring stacking were not the same, and the quaternary fold was completely inverted while the active-site structure remained largely unchanged (Fig.S1). In-depth structural comparison of this prokaryotic GSIII with classes GSI and GSII was thoroughly presented here^23^. Nevertheless, no experimental structural data is available for any eukaryotic GSIII analog.

*Ostreococcus tauri* is the smallest free-living autotrophic eukaryote and an emerging microalgal model organism^24^. Not only is its physical size small, but its genome is also highly compressed with only ∼8000 genes. While most organisms have at least one or more GSI or GSII variants in combination with any GSIII,

*O. tauri*’s annotated genome shows only a single gene with homology to glutamine synthetase (Enzyme Commission Number 6.3.1.2). This gene has two recognized protein domains matching an N-terminal GSIII motif and a C-terminal glutamine synthetase catalytic domain motif. In this report, we identify the native protein sequence of GSIII, its oligomeric functional assembly state and describe its structure, resolved to 3.2Å by cryo-EM and draw comparisons with its prokaryotic structural analogs.

## Materials and Methods

### Culture growth conditions

*Ostreococcus tauri* strain OTTH0595 was obtained from the Roscoff Culture Collection (RCC745). Starter cultures were grown in standard Keller medium^25^ at 22°C with 25 µE/m^2^/sec blue light from 17W T8 50/50 bulbs (Coralife, USA) fitted with 183 Moonlight Blue filters (Lee Filters, USA) on 12-hour light/dark cycles^26,27^. To extract genomic DNA, the cells were harvested from 50 ml of fully grown cultures by centrifugation at 1,000 g for 15 min and subsequently frozen in liquid nitrogen and stored at –80°C. Genomic DNA was extracted using QIAGEN DNeasy Plant Mini Kit as recommended by the manufacturer but omitting tissue homogenization procedures. Genomic DNA was further ethanol precipitated and re-suspended in water.

### The GS gene amplification from genomic DNA, pEU vector construction

Both isoforms of GSIII were amplified through PCR from genomic DNA of *O. tauri*. To amplify the GSIII isoforms, the following PCR primers were used: (1) FWD_longGS: GACGACAAGCTCGCT<u>CAGTCCGTGTTGAGCCA</u>, (2) FWD_shortGS: GACGACAAGCTCG<u>CTCTCTCGATGAGCAGCGG</u>, (3) REV_universal: GCCCTATATATGGATCCTTA<u>GCAGCCGTATTCCGACT</u>. The gene-specific regions are underlined, and the rest of the primer sequences contain the extension regions to create overlaps with the vector template for Gibson cloning^28^. The template vector was pEU_3xflag (modified in house pEU-MCS-TEV-HIS-C1 (CellFree Sciences) to contain a 3xflag tag). All DNA primers were purchased from Invitrogen or IDTDNA. PCR reactions were carried out using the Q5 Hot Start High-Fidelity 2X Master Mix from New England Biolabs. Obtained PCR products, such as linearized pEU template and the gene insert, were gel-purified using Zymoclean Gel DNA recovery kit from Zymo Research and subjected to Gibson Assembly. The Gibson Assembly Master Mix from New England Biolabs was used as recommended. The assembled product was used for *E. coli* transformation, which was later plated on a carbenicillin-containing agar plate and grown at 37°C overnight. Several individual colonies were selected and grown overnight in LB medium, supplemented with carbenicillin. Plasmids were extracted using QIAGEN Plasmid Mini Kit. The insertion of the desired fragment in the plasmid was verified by PCR. The plasmids, that had an insert of the desired size, were sent to MCLAB for sequencing. Successful clones were later grown in larger quantities of LB broth (with carbenicillin) and harvested using the QIAGEN Plasmid Maxi kit.

### Cell-free protein synthesis and purification

Protein synthesis was carried out in the bilayer format using Wheat Germ Protein Research Kit, WEPRO7240, from CellFree Sciences. The detailed protein synthesis and purification procedure was carried out as previously described^29^.

### Negative staining TEM

An ultrathin carbon film on lacey carbon (01824, Ted Pella) was glow discharged at 10 mA for 30 seconds using a PELCO easiGlow from Ted Pella. Then 2 μl of 80 μg/μl GS solution (in TBS buffer) was deposited on the carbon and allowed to absorb for 2 minutes. The excess of solution was quickly blotted away, and the grid was floated on a drop of the NanoW stain from Nanoprobes for 30 seconds. The excess of the stain was wicked away, and the grid was left to air-dry. The TEM micrographs were acquired at -4 μm defocus on a 300kV JEOL JEM-3000SFF using the direct electron detector DE20 from Direct Electron.

### Activity assays

The activity of GSIII was determined by the Mg^2+^- dependent biosynthetic assay, where a released phosphate was detected by the colorimetric Malachite Green Phosphate Assay kit from Cayman Chemical. The standard curve for phosphate standards was generated as instructed by Cayman Chemical. The composition of the Assay Buffer was 25 mM HEPES, pH 7.1, 50 mM NH_4_Cl, 0.4 mM ATP, 3 mM MgCl_2_, 11.5 mM L-glutamate. Four GSIII activity reactions were initiated by mixing 49 μl of Assay Buffer with 1 μl of GSIII (0.05 μg/μl, 0.1 μg/μl, 0.2 μg/μl and 0.4 μg/μl in TBS) and incubated at 37°C for 15 min. The reactions were terminated by addition of 5 μl of MG Acidic Solution from the kit, mixed and incubated for 10 minutes at room temperature. Then 15 μl of MG Blue Solution was supplemented, and incubation continued for extra 20 minutes. The phosphate standards were done in parallel. The absorbance was measured at 620 nm using the Infinite M200 PRO microplate reader by Tecan. All measurements were performed in duplicate. To determine the inhibitory effect of PPT herbicide (from Sigma), the Assay buffer was first supplemented with the desired PPT amount and then combined with 0.4 μg of GS.

### Native Mass Spectrometry (Native MS)

Tagged or untagged GSIII proteins at 3 mg/mL were buffer exchanged into 100 mM ammonium acetate using Zeba Spin Desalting Columns (75 µL, 7K MWCO, ThermoScientific) and further diluted to 0.3 mg/ml in the same ammonium acetate buffer. Mass spectra (*m/z* range from 2500-25000) of intact GS complexes were acquired on a Thermo UHMR Orbitrap operated in positive ion mode. The protein samples were first loaded into pulled glass emitter (outer diameter 1.2 mm, inner diameter, 0.69 mm, tip diameter 2 µm. Part No. BG12692N20, Scientific Instrument Services, Inc., Ringoes, NJ, USA), then a platinum wire at 1 kV was inserted into the emitter to maintain the electrospray. At the ion source, the temperature was set to 200 °C, in source fragmentation at 100 V, s-lens at 50%, and in-source trapping at 200 V. HCD gas was set to 0.75. Resolution was set to 15k where 1000 microscans were averaged. Mass was calibrated externally using cesium iodide clusters. Deconvolution of the raw mass spectra was performed in UniDec^30^.

### Cryo-EM sample preparation, imaging, and single particle analysis

Quantifoil TEM substrates, 658-300-CU from Ted Pella, were first cleaned for 1 hour in chloroform, air dried and then glow-discharged at 15 mA for 60s using PELCO easiGlow. The blotting and vitrification were performed using Vitrobot from FEI. The solution of tag-free GSIII protein (3 μl, 1.2 mg/ml or 0.6 mg/ml, in TBS buffer) was deposited on the grid, blotted for 3s and plunged in the liquid ethane. The images were collected on 300 keV FEI Titan Krios with Gatan K2 direct electron detector. Two datasets were collected: 646 image stacks from the 1.2 mg/ml GSIII stock and 445 image stacks from 0.6 mg/ml GSIII stock. Each image stack had 50 frames at a dosage of 2 e/Å^2^/frame and pixel size of 0.6509 Å. All data processing was performed in cryoSPARC v3.3.1 software^31^. Particle picking yielded an initial pool of 554,543 particles. This particle library was further downsized to 270,133 via three rounds of 2D classification (200 classes, 40 iteration each). These particles were later subjected to Ab-Initio 3D reconstruction with 2 starts (40 cycles per each). The Ab Initio 3D model was further employed as a 3D reference for Auto Refine steps with initial resolution limit of 20Å. All 3D refinements were in C6 symmetry and resulted in 3.18 Å resolution structure.

### Model Building

The initial atomic model of GSIII was generated in Alphafold^32^ and was docked in Chimera^33^. Further atomic model building, and real space refinements were carried out in Coot^34^ and PHENIX^35^ (*Phenix*.*RealspaceRefine*) for an individual monomer. Map symmetry operators were then generated by *Phenix*.*MapSymmetry* and applied to the real space refined monomer model using *Phenix*.*ApplyNCSoperators* to generate the hexamer, which underwent additional rounds of *Phenix*.*RealspaceRefine* (see Table S1 for refinement statistics). The final model and map were uploaded to the PDB and EMDB databases under the accession numbers: **7U6O** and **EMD-26377**.

## Results and Discussion

### Eukaryotic GSIII annotation, its isoforms and proteomics analysis

*Ostreococcus tauri* is believed to have evolved around 1 billion years ago^36^. As such it can provide interesting insight into early evolution of GS enzymes. To date, two different genome annotations identify the same GSIII isoform with the only difference being a potential alternative start codon that would add 35 amino acids on the N-terminus. The newer annotated long form (NCBI ID: CEF96824.1) was the result of a recent de novo resequencing effort in 2014^37^ while the older short form (NCBI ID: XP_003074553.1) was based on shotgun sequencing and assembly performed in 2006^24^.

To determine which annotation of GSIII is natively expressed in *O. tauri*, we reviewed data from our prior global and phosphoproteomic analyses^38^ of wild-type cultures grown in Keller marine media under both normal and nitrogen depleted growth conditions. Under nitrogen deprivation, the GSIII protein was the 10^th^ highest upregulated protein detected in the published datasets, suggesting its potentially critical role in nitrogen metabolism and recycling. Similarly, previous studies done on bacteria also observed an increase in GSIII expression and activity upon nitrogen starvation^39^. From this proteomics data, all LC-MS/MS datasets were screened for peptide coverage against both the long and short GSIII annotations from the published genome assemblies with all detected peptides mapping to the shorter version (Fig.1). Coverage for the detected peptides spanned the protein sequence from the start site for the short annotation to the final arginine, 27 amino acids shy of the C-terminus (representing final primary cleavage site for trypsin). No peptides for the N-terminal sequence of the long version were detected. While this does not prove conclusively that the long version is not natively expressed at all, it is highly suggestive that the predominant functional form is the short version of the protein (XP_003074553.1). To address the questions arising from the above-mentioned observations, we decided to synthesize both (long and short) forms of *O. tauri* GSIII for further biochemical and structural analysis.

**Fig. 1.**
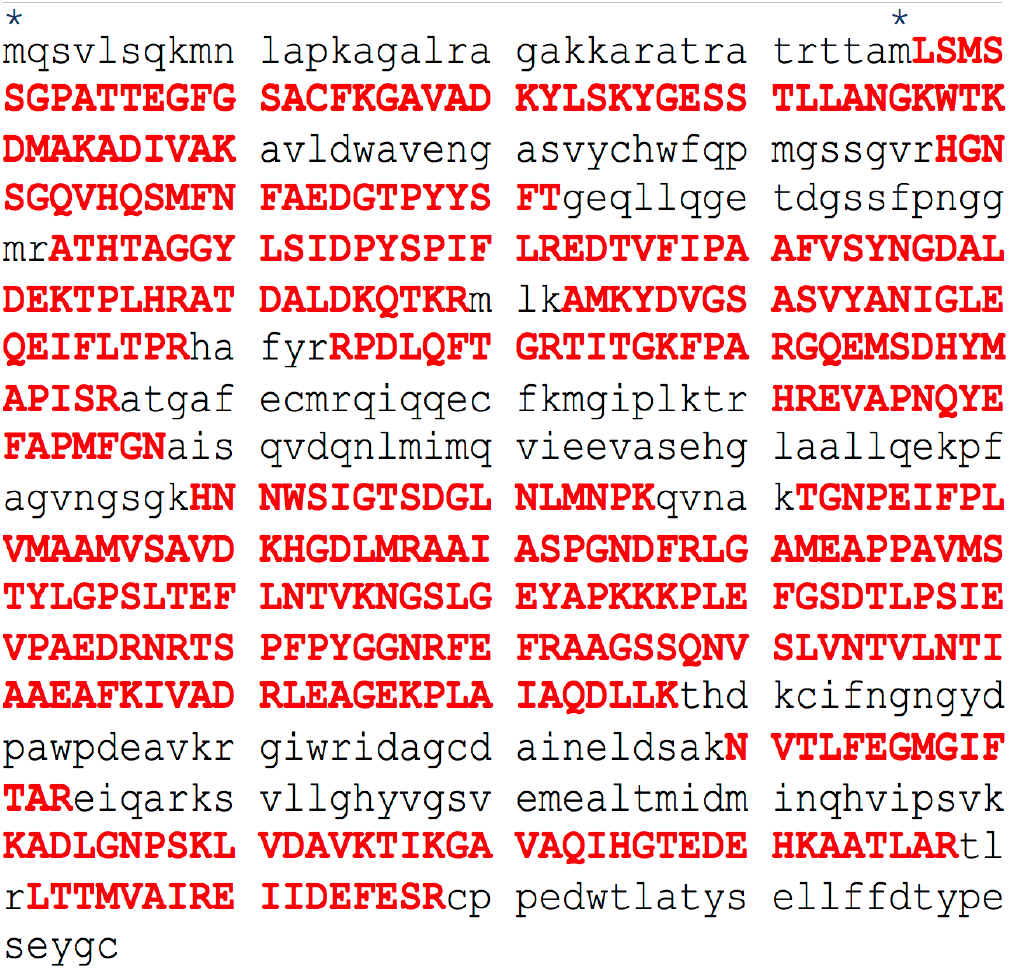
Sequences of both annotations for GSIII from *Ostreococcus tauri* with highlighted LC-MS/MS coverage of peptides detected from lysates of wild-type growth culture. The asterisk symbols indicate the alternate N-terminal methionine start position for the long and short GSIII annotations (left and right asterisks respectively). Detected amino acids are denoted by red capital letters.

### Cell-free expression of GSIII and preliminary characterization by TEM

Both long and short versions of GSIII gene were obtained through PCR amplification from the genomic DNA of *O. tauri* and further cloned into pEU vector, specifically designed for cell-free expression using the wheat germ system^40^. The wheat germ cell-free platform^29,41,42^ was chosen as a good eukaryotic, plant-based system that could provide the closest translational environment for the synthesis of a soluble and functional form of recombinant GSIII. Both short and long isoforms of GSIII protein were well expressed, fully recovered in a soluble fraction and were efficiently purified using a cleavable 3xFLAG tag (Fig.2a). The theoretical molecular weights of short and long GSs were approximately 80 kDa, and both species ran on denaturing polyacrylamide gel with expected mobility. Cell-free produced and purified GSIII long and short forms also appeared to be full-size, homogenous, and free of any premature termination products with purity above 98%. As the last stage of protein synthesis and purification, the 3xFLAG tag was removed with enterokinase.

Obtained short and long isoforms of GSIII protein were further subjected to preliminary analysis by NanoW negative stain TEM imaging. Electron micrographs (Fig. 2b,2c) show a dense protein complex of pinwheel architecture with an average diameter of 16.5 nm for both isoforms. In the case of short GSIII, few aggregation events or partially disordered complexes were seen, whereas the long GSIII isoform exhibited a higher proportion of smaller assemblies and disordered complexes as well as a larger central pore to the pinwheel. We posit these changes seen for the long isoform are most likely due to the extra 35 amino acids at the N-terminus that can cause the perturbation in complex stability or assembly. Due to lower sample homogeneity of the long GSIII isoform and the lack of the support for its existence by proteomics, all subsequent experiments were conducted on the short GS variant.

**Fig. 2.**
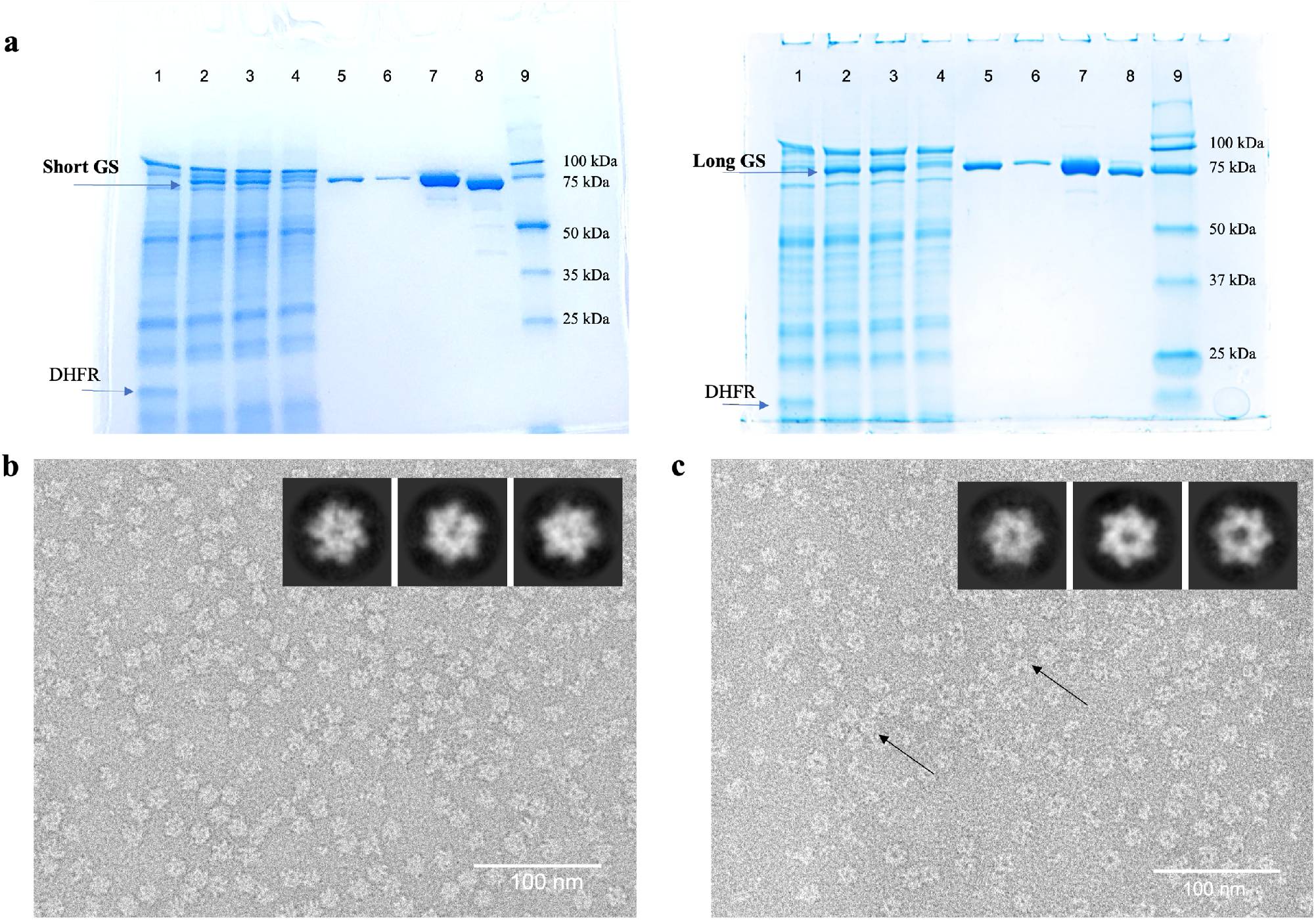
GSIII production and characterization. a. SDS-PAGE gels showing expression and purification of short GSIII (on the left) and the long GSIII (on the right). Lane 1, control DHFR expression; Lane 2, crude mixture; Lane 3, soluble fraction; Lane 4, flow-through; Lane 5, elution fraction 1; Lane 6, elution fraction 2; Lane 7, elution fractions 1 and 2 combined and concentrated; Lane 8, final protein with 3XFLAG cleaved off; Lane 9, MW markers. b. Negative stain TEM imaging of short GSIII. c. Negative stain TEM imaging of long GSIII. Black arrows point to the areas of disordered protein complexes.

### *O. tauri* GSIII exists as a defined and stable 6-mer oligomer of 463 kDa and is functional

The only high-resolution structure of a type III GS available today is from the bacterial organism *Bacteroides fragilis*^23^. Based on that prokaryotic 3D architecture, GSIII complexes are expected to exist as dodecameric two-ring self-assembly with a 6-fold dihedral space group symmetry. However, sequence alignment of *O. tauri* GSIII against *B. fragilis* GSIII shows less than 50% homology, 30% identity and 10% gaps (Fig.S1). To evaluate the *O. tauri* GSIII oligomerization, we analyzed cell-free expressed complexes with Native PAGE in the absence and in the presence of Mg^2+^ and Mn^2+^ ions since these cations have both been previously reported to induce and stabilize stacked ring assembly^43^ (Fig.3a). In every condition, *O. tauri* GSIII migrated as a single well-defined complex with an approximate molecular weight of 500-550 kDa, which is in the range of a 6mer or 7mer complex. While it is known that proteins with Native PAGE can run faster or slower than predicted based on their shape or charge, the lack of any higher mass band under any condition tested provided initial evidence that this eukaryotic GSIII differs in functional assembly state compared to the 12-mer complex reported for *B. fragilis* prokaryotic GSIII. Interestingly, a publication from 1997 of a cyanobacterial prokaryotic GSIII suggested a functional hexamer may exist^44^ and an early report of *B. fragilis* GSIII from 1987 suggested it also existed as a hexamer^45^ all based on biochemical data before later cryo-EM experiments showed native assembly and function as a dodecamer.

To obtain an accurate molecular weight of *O. tauri* GSIII and unambiguously define subunit stoichiometry we used native mass spectrometry (native MS) (Fig. 3b,c). The mass spectrum of GSIII electrosprayed from 100 mM ammonium acetate solution (pH 7) predominantly showed the signal from a 463.4k species, corresponding to a hexamer of GSIII (77156.9 Da average theoretical mass of tag-removed 1mer, corresponding to 462.9k 6mer mass). The slightly higher experimental mass can be explained by noncovalent binding of salts/solvents and/or some modifications (oxidation, N-terminal acetylation). We also detected peptides containing part of the N-terminal tag (mass of full tag is 2.8 kDa) after digesting the protein with trypsin, indicating incomplete tag removal (Fig. S2). Those species at 465.0k, 466.3k, and 467.8k were assigned to be GSIII 6mers containing 1mer(s) with incomplete tag cleavage in a small population. Without tag removal, the GSIII protein was detected as a 6mer (Fig. S2) at 480.4 kDa. The 6mer is very stable in the gas phase and did not dissociate or release 1mers even at the maximum collision energy, indicating *O. tauri* GSIII 6mer has strong inter-subunit interfaces. No other species beyond the 6mer were observed.

**Fig. 3.**
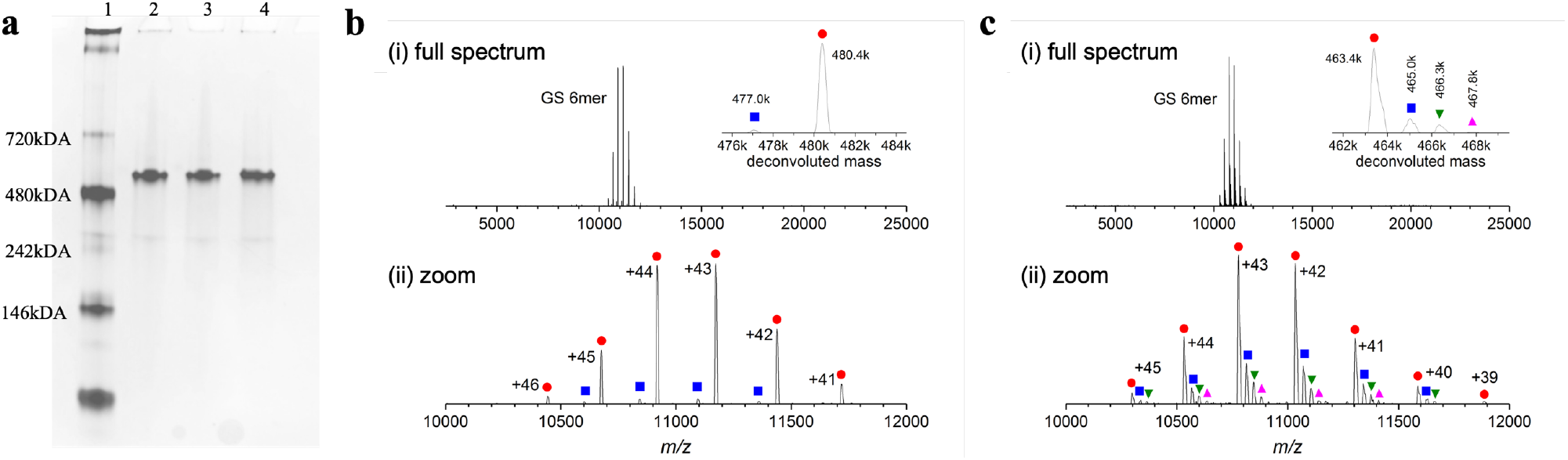
The folding of GSIII. a. Native PAGE analysis. Lane 1, NativeMark Protein Standard; Lane 2, GSIII in TBS buffer; Lane 3, GSIII in TBS buffer supplemented with 10 mM MgCl_2_; Lane 4, GSIII in TBS buffer supplemented with 10 mM MnCl_2_. b. Native mass spectra of the 3XFLAG-tagged eu-GSIII protein shown in (i) full mass range, and (ii) zoomed view of the 6mer region. Inset plot in (i) shows the deconvoluted mass distribution of tagged GS 6-mer. Each species at a particular mass displays several charge states in the raw mass spectrum. The charge states are annotated in (ii) next to each peak cluster. c. Native mass spectra of the tag-free eu-GSIII shown in (i) full mass range, and (ii) zoomed view of the 6mer region. Inset plot in (i) shows the 6mer deconvoluted mass distribution, with masses listed next to the peaks. The peaks with mass larger than 463.4k originated from GSIII 6mers containing 1mer(s) with incomplete tag cleavage. The same symbols are used in the inset and (ii).

All forms of GS use ammonium ions and glutamate as substrates in ATP-dependent reactions to synthesize glutamine^46^. Thus, to determine if the obtained stable hexameric GSIII complex was functional, the biosynthetic activity of the protein was assayed by detecting the amount of Pi released during this reaction using a colorimetric approach. The assay buffer and incubation conditions were chosen from the previous study^47^. The amount of released phosphate by GSIII has been interpolated on the standard phosphate graph (Fig. S3). Through the linear fit of GSIII amounts versus released phosphate, the specific activity of GS at these assay conditions and with saturating amounts of substrate was determined to be 0.13 nmoles/min/μg protein. As discussed previously, GS proteins are the target of irreversible inhibition by PPT herbicide^1^, therefore, to further validate complex functionality, we assayed the activity of the enzyme in the presence of varying amounts of PPT (Fig. S3c). As expected, *O. tauri* GSIII activity was dramatically affected in the presence of PPT, a reduction of 50% observed at 0.4 mM PPT and only 20% of its activity remained at 2 mM PPT.

### The 3.2Å cryo-EM structure of GSIII

We employed single particle cryo-EM to determine the atomic resolution structure of *O. tauri* GSIII since prior to the present study, no experimentally validated structure of a eukaryotic GSIII exists. Acquired micrographs showed individual GSIII hexamers at different orientations (Fig. S4). The raw images and 2D class averages confirm the single stacked hexameric pinwheel morphology seen with TEM negative staining earlier. The reconstructed 3D volume exhibits C6 symmetry and was resolved to 3.18Å resolution. The local resolution colored plot overlaid on the refined map (Fig. S4f) depicts resolution ranging from 2.8Å to 4.4Å with the lower resolution features found at the periphery of the complex. We tested local 3D refinements using masks for monomers, dimers, trimers, and the whole hexamer to evaluate if there was intramolecular asymmetry that would contribute to the lower resolution at the periphery. All of these refinement trials showed similar FSC convergence and monomer density maps suggesting there is just naturally more flexibility near the complex periphery.

To interpret the 3D cryo-EM map, we used an Alphafold2-generated model of *O. tauri* GSIII as an initial model for model building and refinement. The model docked fairly well but required multiple rounds of manual adjustment in COOT and automated fitting with *Phenix*.*RealSpaceRefine* to adjust the alpha carbon trace and side chains to fit the experimental density. No associated electron density for N-terminal residues 1-37 and C-terminal residues 705-729, containing purification tags, were present in the final 3D map suggesting the lack of defined folds which is generally somewhat expected for terminal ends. Select regions of 3D volume, which correspond to the surface exposed internal loops, showed an electron density but of low quality. The amino acids (Val105-Gly108, Phe145-Gly158 and Met264-Thr276) from these lowquality regions have been removed from the final atomic model since their correlation values did not exceed Interestingly, the crystal structure of prokaryotic GSIII (PDB ID:3O6X) also failed to resolve those equivalent residues suggesting an inherent flexibility of these regions in the GSIII class as a whole.

### The structural comparison between eu-GSIII and pro-GSIII

A very thorough structural comparison between GS classes I, II and the single structure of prokaryotic GSIII have been reported previously^23^. Therefore, all further comparisons discussed below will focus only on difference between the 2 experimentally determined GSIII from the prokaryote *B. fragilis* (pro-GSIII) and the eukaryote *O. tauri* (eu-GSIII), the structure of which is presented here (Fig. 4). Secondary structure elements are named as specified herein (Fig.S6). Previously, it was shown that pro-GSIII forms a dodecamer with 622 (D6) symmetry in solution by cryo-EM^20^, and its high-resolution crystal structure^23^ was refined for a hexameric ring as the asymmetric unit with a unit cell symmetry of C222_1_ that recreates the dodecamer. Unlike pro-GSIII, eu-GSIII exclusively forms hexamers in solution both structurally and biochemically. The atomic structure of eu-GSIII exhibits similarities in the overall shape to the hexameric ring of pro-GSIII (Fig.4a). Noticeable differences manifest in the areas surrounding the central opening and at the periphery of the complex. The central cavity of pro-GSIII complex is narrower because of dense structural arrangements of **α** helices. At the periphery, in eu-GSIII, beta strands appear to extend the interaction interface between monomers to form a clasp. Those differential features are best distinguished by looking at a monomeric unit first (Fig.4b). Specifically, helices **α13** and **α14** are significantly shortened in eu-GSIII. The latter widens the central tunnel in eu-GSIII. The helix **αA** of pro-GSIII does not exist in eu-GSIII due to a truncation of the N-terminus. An internal loop (residues Gly446-Asp474), not structurally resolved in pro-GSIII, shows the formation of two beta sheets **βC’** and **βC’**’. Additionally, **β2** is better resolved in eu-GSIII, which is a conserved secondary element in other GSI and GSII classes.

**Fig. 4.**
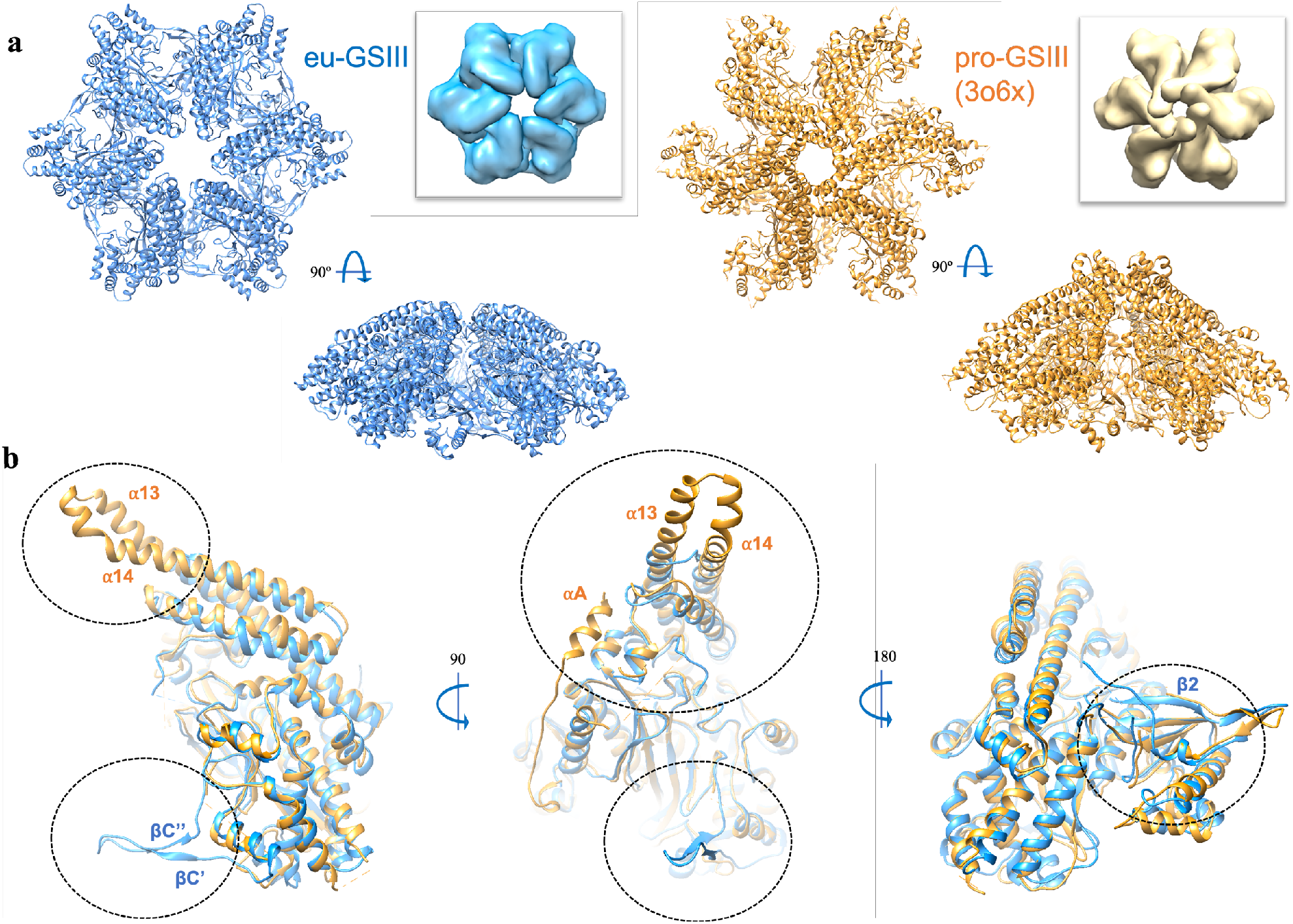
The atomic structure of GSIII hexamer from *O. tauri* (eu-GSIII) compared with PDB 3O6X from *B. fragilis* a prokaryotic homolog (pro-GSIII). a. Top and side views of atomic models for eu-GSIII (in blue) and pro-GSIII (in beige). Inset images depict surface models to highlight gross topological differences. b. Various side-views of superimposed eu-GSIII and pro-GSIII monomers. The regions of significant variance between the two proteins are shown in dashed circles. RMSD between 432 pruned atom pairs is 1.113 Å (across all 567 pairs is 3.252 Å). The secondary structure annotations are as specified in Fig. S6.

Each hexamer contains 6 sites for catalysis, each located at the interface of individual subunits, between the N-terminal region of one monomer and several β-strands of the C-terminal of an adjacent subunit. Superimposed images of the catalytic sites for eu-GSIII and pro-GSIII demonstrate clear conservation of the catalytic core (Fig.S7). The N-terminal region containing key residues Asp141 and Ser144 in eu-GSIII (which correspond to Asp129 and Ser132 in pro-GSIII) have been resolved in eu-GSIII, most likely, due to stabilizing neighboring interaction of **βC’** and **βC’**’ with helix **α3**.

The main deviations are observed in the quaternary structure (Fig. 5). It has been determined already that quaternary contacts are formed between the least conserved amino acid regions and show significant deviation from class to class^23^. The protein-protein interface analysis of eu-GSIII showed that the interface area is close to the value of ∼1930Å^2^, which is slightly weaker than 2262 Å^2^ in pro-GSIII (Fig.5a) but it is still quite notable and higher or equal to previously reported values of ∼1552Å^2^ and ∼1890Å^2^ for intra-ring interfaces for GSI and GSII respectively^23^.

**Fig. 5.**
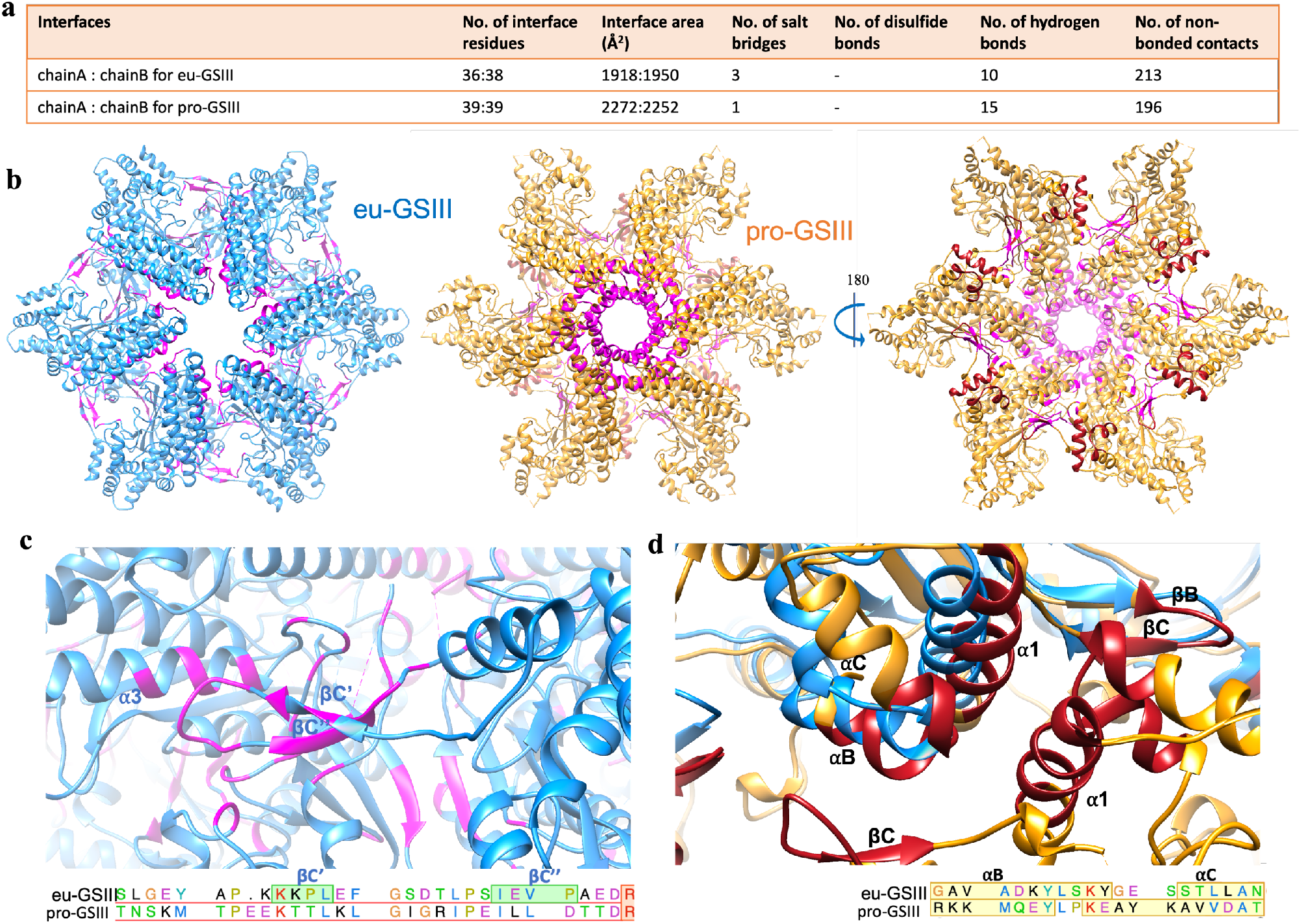
Quaternary structure analysis of GSIIIs. a. The interface values (from PDBsum analysis) are presented between chains A and B only, as all six chains A, B, C, D, E and F are equivalent. b. Interface residues for the hexamer intra-ring formation are mapped in magenta. Interface residues for the inter-ring formation of dodecamer in pro-GSIII are shown in red. c. A close-up view of **βC’** and **βC’’** sheets of chain A interacting with **α3** of chain F. d. A close-up view of inter-ring dodecamer interface formed in pro-GSIII. The hexamer eu-GSIII (in blue) is only superimposed with one hexamer from pro-GSIII.

The residues involved in the formation of the intra-ring contacts show distinct patterns between the two GSIII structures (Fig.5b). Unlike pro-GSIII, the core intra-ring contacts of eu-GSIII are not centered around the central pore of the complex and rather distributed alongside the subunit-subunit interface. A new beta hairpin protein motif, comprising of **βC’** and **βC’**’ strands, interacts with an internal loop and helix **α3** of the adjacent subunit, locking the hexamer fold on the periphery (Fig.5c). The site, where inter-rings contacts are formed to shape a dodecamer in pro-GSIII, differs significantly in the positional orientation of the helix **αC** (Fig.5d). Helices **αB** and **α1** of eu-GSIII are shifted slightly inward. Unfortunately, the resolution limits of both structures do not allow reliable in-depth comparison of amino acid side chains at these sites.

## Conclusion

In this work, we characterized and presented the first structure of eukaryotic type III glutamine synthetase to expand knowledge of this important family of proteins. We identified that eu-GSIIIs fold and act exclusively as stable hexamers. The catalytic core and many other secondary and tertiary elements at a monomer level have been shown to be well-conserved in the GS family, and the structure of eu-GSIII did not deviate from the above. However, the catalytic site is positioned alongside quaternary interfaces that can impact drug/inhibitor binding. Key variations between GS classes are observed in those quaternary interactions in both intra-ring and inter-ring contacts. Intra-ring interactions are weakened in the center of the ring for the *O. tauri* eu-GSIII hexamer, but include beta-hairpins, which are new peripheral quaternary stabilization motifs for the GS family that have not been observed in GSIII nor in other classes of the GS family (Fig. 5c). Notably, beta-hairpin motifs in proteins are generally one of the simplest, most commonly occurring, and the most stable motif with a broad functional repertoire, including protein-protein binding^48^. Considering that *O. tauri* evolved 1 billion years ago, it is not surprising that the simplified hairpin is a motif of choice in *O. tauri* GSIII lacking ring-ring stacking. While most GS proteins form primarily 2-ring-stacked complexes, the inter-ring stacking does not happen using the same interfaces between classes. Specifically, hexameric rings in pro-GSIII stack via an opposite surface relative to the GSIs and GSIIs^23^. Additionally, it has been implied that this stacking does not play a role in GS regulation based on a significant variation in its associating interfaces^23^. The structure of eu-GSIII presented here provides further support to that hypothesis since the ring-ring interactions are absent in the single ring hexamer. Furthermore, the observed lack of stacking did not impact functional performance in eu-GSIII, suggesting that this assembly might be beneficial for some GSs but not essential. In addition, the structure of *P. aeruginosa* PA5508 (a homologue of glutamine synthetase) contains a unique folding of C-terminus that aborts any potential ring-ring stacking^49^ and exists as a functional hexamer. Although the PA5508 enzyme has evolved to use different substrates (complex amines instead of ammonia) to synthesize gamma-glutamyl derivatives (instead of glutamine), it is noteworthy to mention that ring-stacking quaternary interaction was not preserved. For GSIII, there is a significant lack of experimental structural or biochemical data in general, however, a report from 1997 on a cyanobacterial GSIII highlights biochemical data that it folds and functions purely as a homo-hexameric complex as well^44^, supporting the idea that ring-stacking is not essential and required feature for type III GS complexes. As GSs are essential proteins for nitrogen metabolism in any organism and are key targets for drug design, understanding their biochemistry, structural similarities and/or differences are of paramount importance, this report expands our knowledge in uncovering their structural diversity.

## Acknowledgements

This research was performed using the Environmental Molecular Sciences Laboratory (EMSL), a national scientific user facility sponsored by the Department of Energy’s office of Biological and Environmental Research and located at Pacific Northwest National Laboratory (PNNL). This work was supported by DOE-BER FWPs #66382 and 74915.

