## Supplemental Materials for "Structure Of Eukaryotic Type III Glutamine Synthetase Lacking Ring-Ring Quaternary Contacts"

**Supplementary Materials**

**Peptide Mapping.** 2 µL (6 µg) of the buffer exchanged GS proteins used for native MS were diluted to 10 µL in water and mixed with 10 µL trifluoroethanol. The mixture was further incubated in the thermomixer at 60 ºC and 800 rpm for 2 hours. Then 5mM dithiothreitol was added and incubated at 50 ºC and 800 rpm for additional 1 hour. After that, 200 µL of 25 mM ammonium bicarbonate and 0.1 µg trypsin were added to digest the protein at these conditions: 37 ºC, 800 rpm, overnight. The sample was then dried in SpeedVac and re-constituted to 0.05 µg/µL in 0.1% formic acid. Peptides were loaded on to a Waters NanoAcquity liquid chromatography system with C18 reversed phase column (packed in-house, length 70 cm, 75 µm ID, 3µm Jupiter particle from Phenomenex). Peptides were separated using water/acetonitrile mobile phases with 0.1% formic acid over a gradient of 100min (ramping acetonitrile gradient 1-40%) at a flow rate of 300 nL/min. Mass spectra of eluting peptides were collected on a Thermo Lumos Orbitrap. Resolution of 60k and 50k (at *m/z* 200) were used for $MS^1$ and $MS^2$, respectively. A cycle time of 2 s was used to select precursors (with charge 2-10) for higher energy collision (HCD) at 32% normalized collision energy. A dynamic exclusion of 45 s was used. LCMS data were analyzed with Byonic (ProteinMetrics, Cupertino, CA, USA) with semi-tryptic search against wheat protein database (UniProt), including common contaminants, and appended with GS protein sequence including the purification tag. Precursor and fragment mass tolerances were set to 7 and 5 ppm. Protein N-terminal acetylation and

```
CLUSTAL O(1.2.4) multiple sequence alignment

O. tauri    MQSVLSQKMNLAPKAGALRAGAKKARATRATRTTAMLSM-SSGPATTEGFGSACFKGAVA 59
B. fragilis -----MSKMRFFA---------LQELS---NR--KPLEITTPSNKLSDYYASHVFDRKKM 41
                  .**.:            :  :   .*    *.: : .   :: :.*  *.

O. tauri    DKYLSKYGESSTLLANGK-WTKDMAKADIVAKAVLDWAVENGASVYCHWFQPMGSSGVRH 118
B. fragilis QEYLPKEAYKAVVDATEKGTPISREMADLIANGMKSWAKSLNVTHYTHWFQPLTDGT--- 98
            ::** * . .:.: *. *    .   **::*:.: .** . ..: * *****: ..

O. tauri    GNSGQVHQSMFNFAEDGTPYYSFTGEQLLQGETDGSSFPNGGMRATHTAGGYLSIDPYSP 178
B. fragilis ---AEKHDGFIEFGEDGEVIERFSGKLLIQQEPDASSFPNGGIRNTFEARGYTAWDVSSP 155
               .: *:.:::*.***     *:*: *:* * *.*******:* *. * ** : *  **

O. tauri    IFLREDTVFIPAAFVSYNGDALDEKTPLHRATDALDKQTKRMLKAMKYDVGSASVYANIG 238
B. fragilis AFVVDTTLCIPTIFISYTGEALDYKTPLLKALAAVDKAATEVCQ--LFDKNITRVFTNLG 213
             *: : *: **: *:**.*:*** **** :*  *:** :..: :   :* . : *::*:*

O. tauri    LEQEIFLTPRHAFYRRPDLQFTGRTITGKFPARGQEMSDHYMAPISRATGAFECMRQIQQ 298
B. fragilis WEQEYFLVDTSLYNARPDLRLTGRTLMGHSSAKDQQLEDHYFGSIPPRVTA--FMKELEI 271
             *** **.    :  ****::****: *:  *:.*::.***:. *   . *   *::::

O. tauri    ECFKMGIPLKTRHREVAPNQYEFAPMFGNAISQVDQNLMIMQVIEEVASEHGLAALLQEK 358
B. fragilis ECHKLGIPVKTRHNEVAPNQFELAPIFENCNLANDHNQLVMDLMKRIARKHHFAVLFHEK 331
            **.*:***:****.******:*:**:* *.    *:* ::*::::.:* :* :*.*::**

O. tauri    PFAGVNGSGKHNNWSIGTSDGLNLMNPKQVNAKTGNPEIFPLVMAAMVSAVDKHGDLMRA 418
B. fragilis PYNGVNGSGKHNNWSLCTDTGINLFAPGKNPK--G-NMLFLTFLVNVLMMVHKNQDLLRA 388
            *: ************: *. *:**: * :     *   :*  .:. ::  *.*: **:**

O. tauri    AIASPGNDFRLGAMEAPPAVMSTYLGPSLTEFLNTVKNGSLG-E-YAPKKKPLEFGSDTL 476
B. fragilis SIMSAGNSHRLGANEAPPAILSIFLGSQLSATLDEIVRQVTNSKMTPEEKTTLKLGIGRI 448
            :* * **..**** *****::* :** .*:  *: : .   . :    :*. *::* . :

O. tauri    PSIEVPAEDRNRTSPFPYGGNRFEFRAAGSSQNVSLVNTVLNTIAAE--------AFKIV 528
B. fragilis PEILLDTTDRNRTSPFAFTGNRFEFRAAGSSANCAAAMIAINAAMANQLNEFKASVDKLM 508
            *.* : : ******** : ************ * : .  .:*:  *:        . *::

O. tauri    ADRLEAGEKPLAIAQDLLKTHDKCIFNGNGYDPAWPDEAVKRGIWRIDAGCDAINELDSA 588
B. fragilis EEGIGKDEAIFRILKENIIASEPIRFEGDGYSEEWKQEAARRGLTNICHVPEALMHYMDN 568
             : :  .*  : * :: : : :   *:*:**.  * :**.:**: .*    :*: .  .

O. tauri    KNVTLFEGMGIFTAREIQARKSVLLGHYVGSVEMEALTMIDMINQHVIPSVKKAD---LG 645
B. fragilis QSRAVLIGERIFNETELACRLEVELEKYTMKVQIESRVLGDLAINHIVPIAVSYQNRLLE 628
            :. ::: *  **.  *: .* .* * :*. .*::*: .: *: :*::* . . :   *

O. tauri    N--------------------------PSKLVDAVKTIKGAVAQ---IHGTEDEHKAATL 676
B. fragilis NLCRMKEIFSEEEYEVMSADRKELIKEISHRVSAIKVLVRDMTEARKVANHKENFKEKAF 688
            *                           *: *.*:*.:   :::   : . :::.*  ::

O. tauri    A-RTLRLTTMVAIREIIDEFESRCPPEDWTLATYSELLFFDTYPESEYGC 725
Q5LGP1      AYEETVRPYLESIRDHIDHLEMEIDDEIWPLPKYRELLFTK--------- 729
            * .      : :**: **.:* .   * * * .* **** .
```

Fig.S1. Clustal sequence alignment of eukaryotic GSIII from *O. tauri* versus prokaryotic GSIII from *B. fragilis.* The yellow highlight indicates the extra amino acids that would exist due to alternate N-terminal methionine start position for the long versus short GSIII annotation.

(a) Coverage map for tagged GSIII

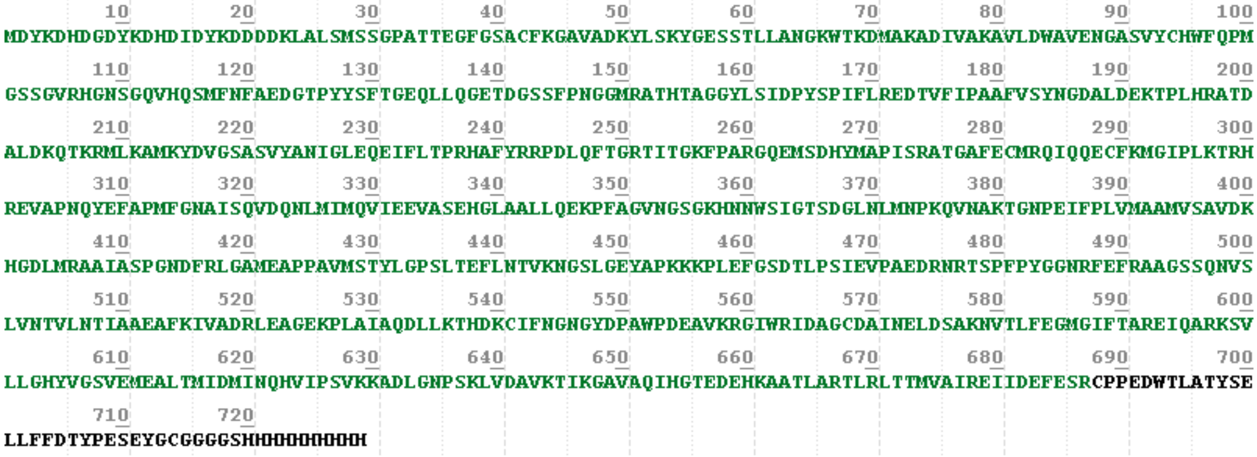

(b) Coverage map for tag-removed GSIII

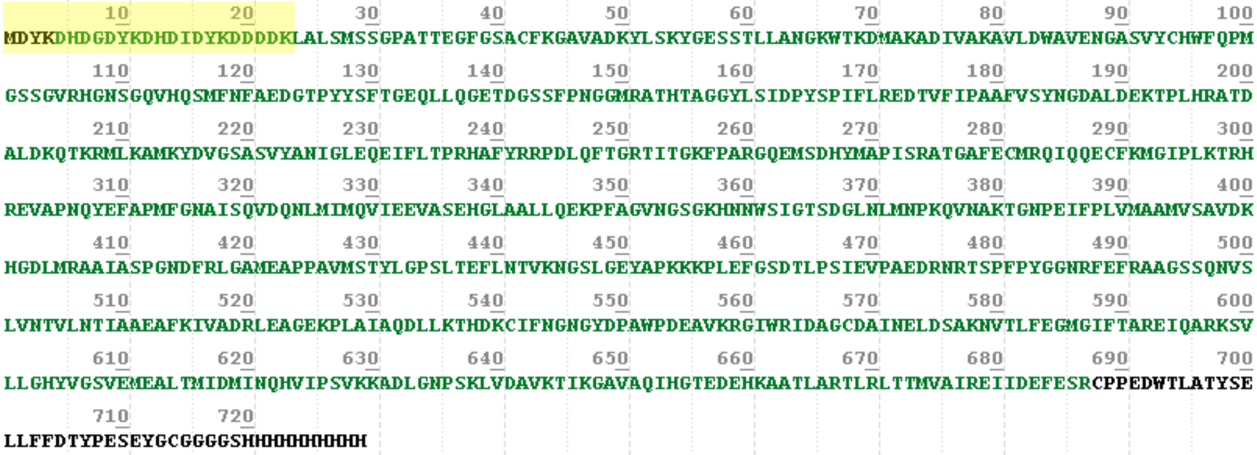

(c) Peptide map for the N-terminal region of the tag-removed GSIII

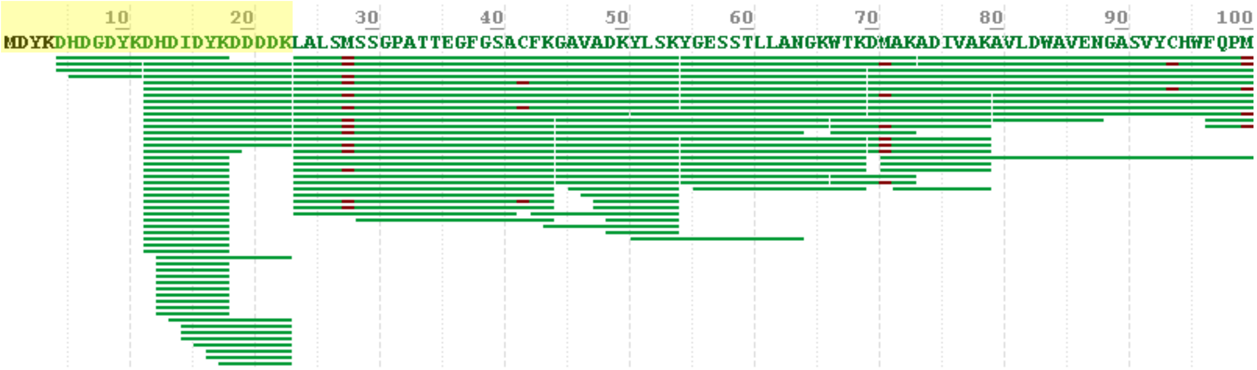

Figure S2. Peptide mapping result showing the coverage of the (a) tagged and (b) tag-removed GSIII samples where amino acids shown in green are detected and amino acids in black are not. Peptide map for the first 100 residues in the sequence for the tag-removed GSIII sample is shown in (c). Each green line below the sequence indicates a peptide was identified within that region. M and C highlighted in purple are sites identified with oxidation. We detected peptides within the tag region (first 23 residues highlighted in yellow background) in (b-c) even after tag removal. This incomplete cleavage by the enzyme gave rise to the additional peaks observed in the native MS spectra. The C-terminal peptide was not detected likely due to its unique property

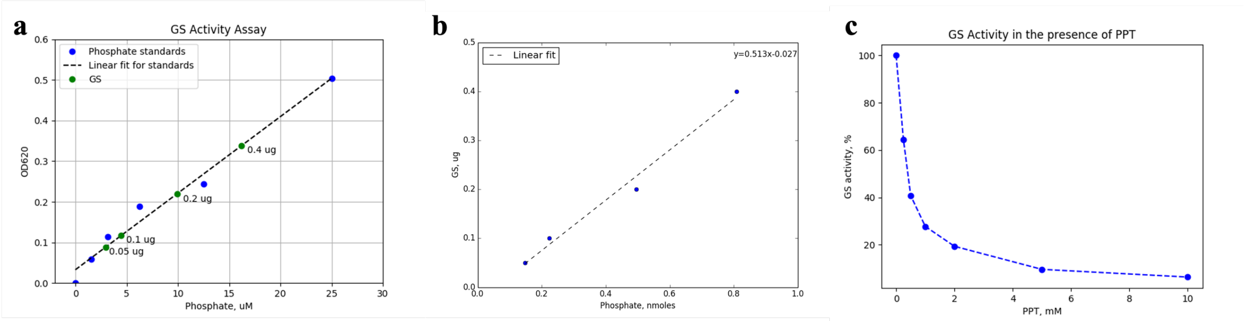


Fig. S3. Functional assays. a. Standard phosphate curve obtained using colorimetry with interpolated phosphate release amounts by GSIII. b. In 50 µl reaction volume, the plot shows the amount of phosphate released (nmoles) depending on the amount of GSIII (µg) introduced in the assay. Through a linear fit, it was determined that the specific activity of GSIII was 2 nmoles/µg at these assay conditions (15 min incubation period). c. The plot of GSIII activity in the presence of various amounts of PPT inhibitor.

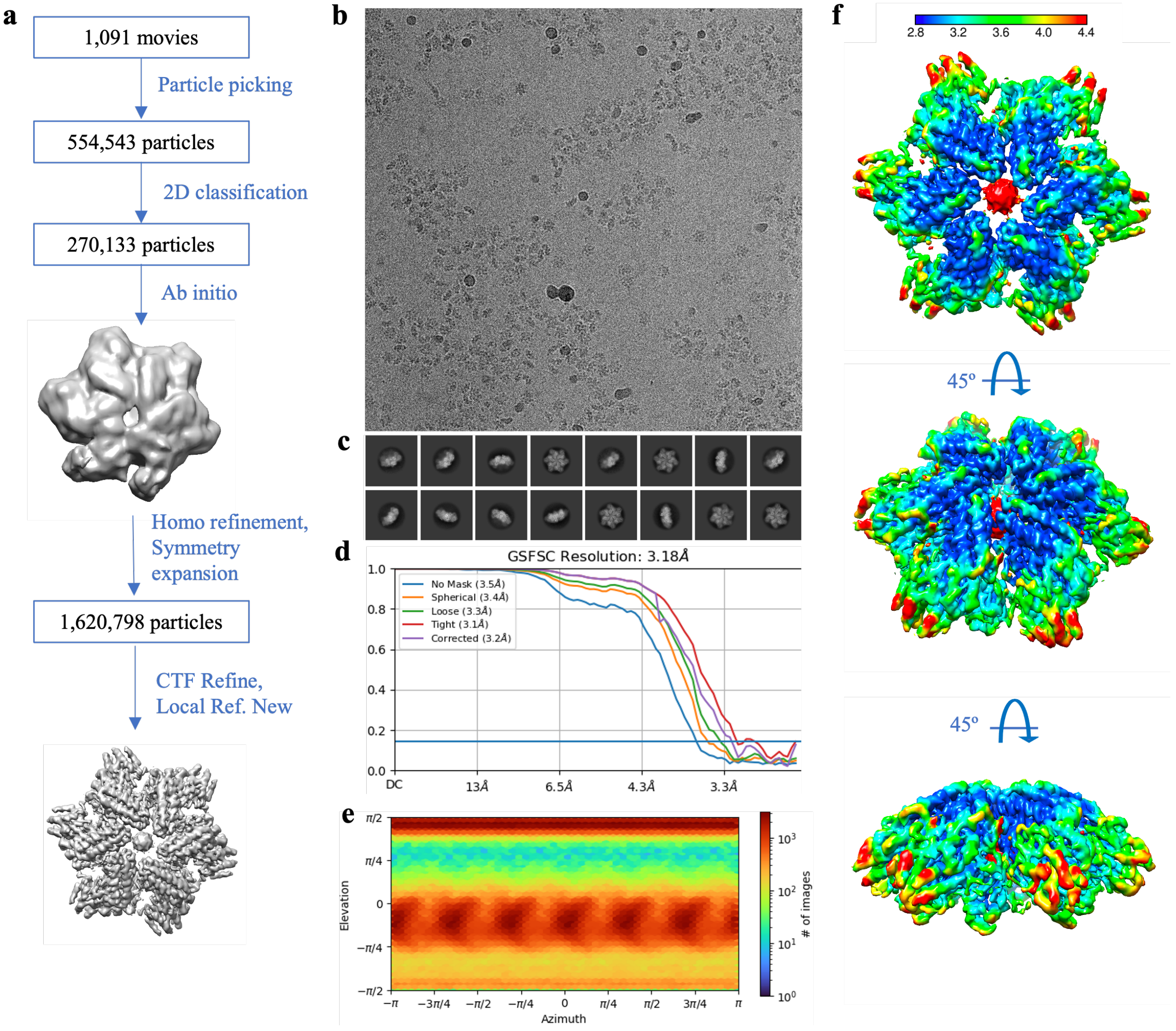


Fig. S4. Cryo-EM single particle analysis workflow for eu-GSIII hexamer from *Ostreococcus tauri*. **a**, All processing was performed in cryoSPARC v3. The positions of particles were identified using a template-based autopicking algorithm. The final pool of 554,543 particles was subjected to several rounds of reference-free 2D classification (with 150 classes each round). The curated pool of 270,133 particles was then used for *ab initio* model generation, which was further used for homogenous refinement. The particles were further symmetry expanded and used in a local refinement to generate a final map of 3.18 Å resolution. **b**, A representative image of the GSIII complexes in vitreous ice. **c**, Representative 2D classes. **d**, The Fourier shell correlation (FSC) curve and **(e)** Euler angle distribution of refined map. **f**, GSIII reconstructed map colored according to local resolution in Angstroms.

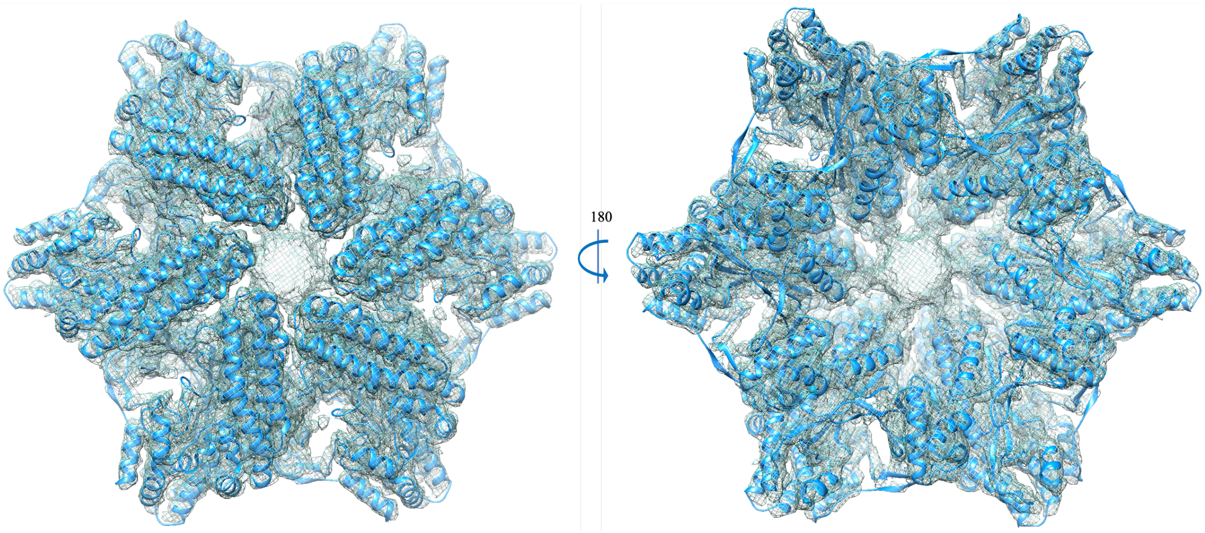


Fig. S5. Atomic model of eu-GSIII fitted in the unsharpened cryo-EM map (grey mesh).

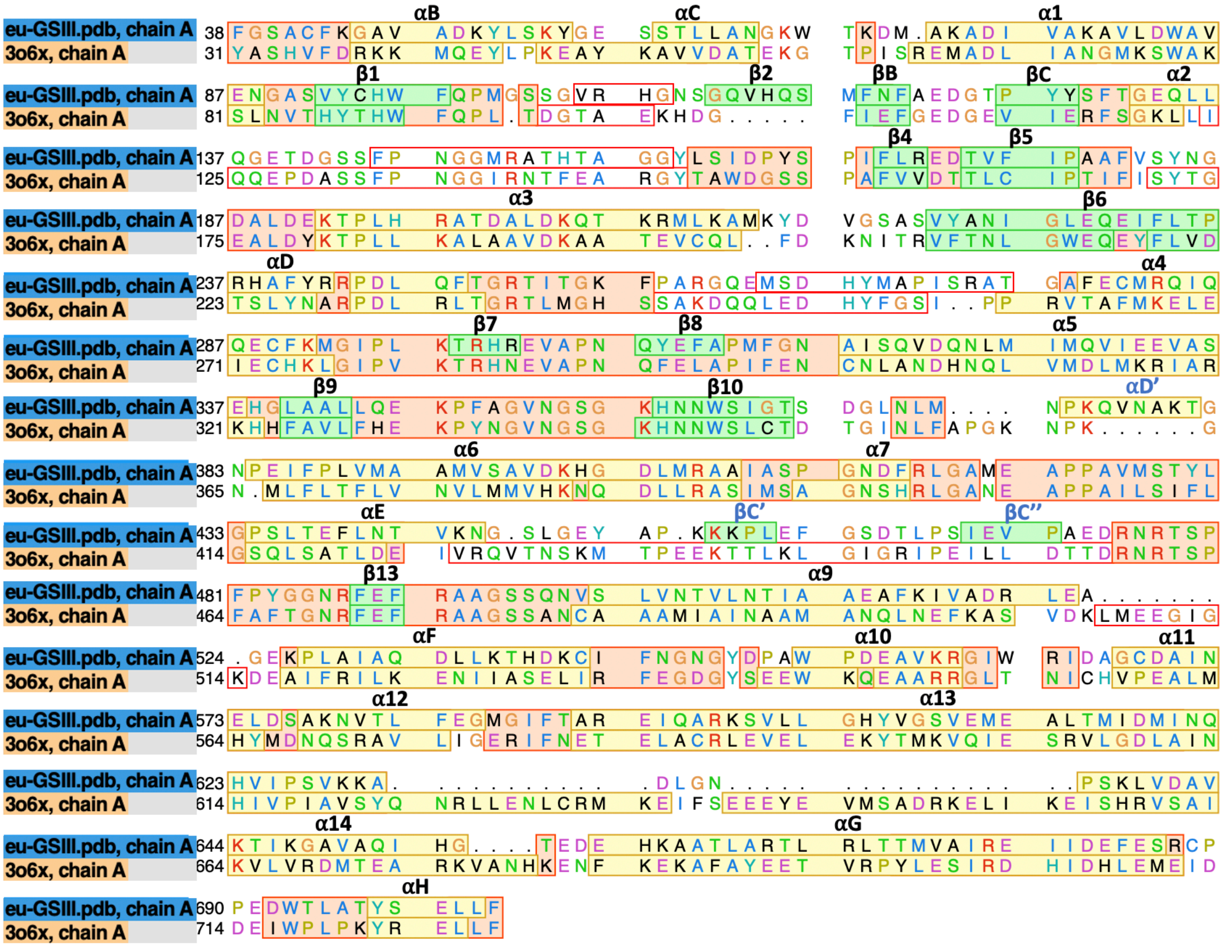

Fig. S6. Structure-based sequence alignment between *Ostreococcus tauri* eu-GSIII and the pro-GSIII crystal structure (3O6X) from *Bacteroides fragilis*, performed in Chimera. The residue coloring is in Clustal X default format. α-Helices (yellow) and ß-sheets (green) are annotated as herein[23]. Matched residues are in light pink; regions depicting missing structure are boxed in solid red. New secondary structure elements observed in eu-GSIII, but not previously annotated in pro-GSIII, are denoted as αD', βC' and βC''.

**a** **b**

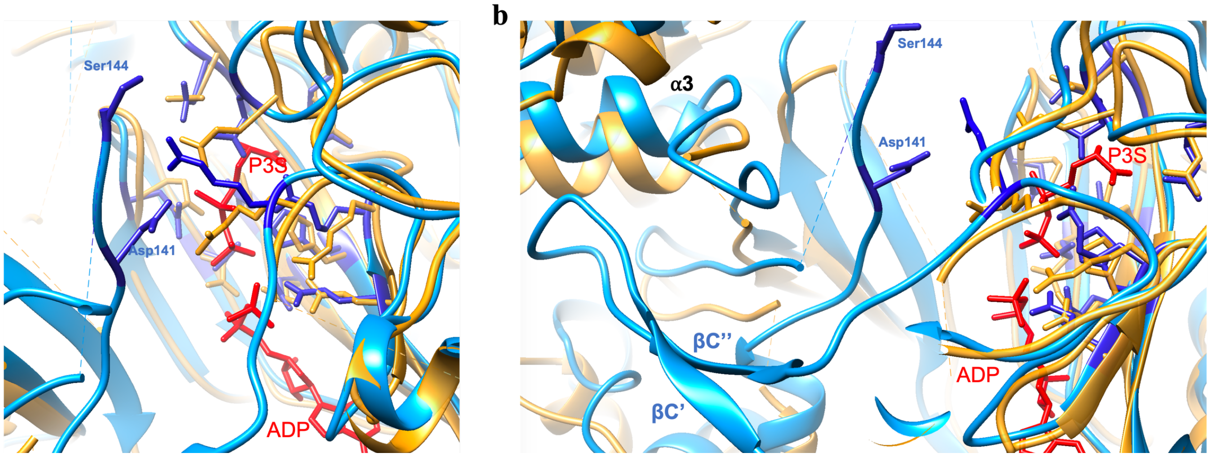


Fig. S7. The close-up views (a) and (b) of the superimposed catalytic site from different angles. The acquired atomic structure of eu-GSIII is in blue. The crystal structure of pro-GSIII and precursor molecules (ADP and P3S) are in beige and red respectively (PDB ID: 3o6x). Conserved active site residues across all GS classes[1] are accentuated in medium blue color on eu-GSIII (Asp141, Ser144, Glu229, Glu231, Glu302, Glu309, Asn353, Gly354, His358, Arg417, Glu422, Arg477, Glu489, Arg491). We also show the side chains of the same residues on pro-GSIII for comparison. Note that the prior published crystal structure for pro-GSIII did not resolve residues Asp129 and Ser132 in the active site and are therefore not shown whereas the corresponding residues in eu-GSIII (Asp141 and Ser144) are resolved in the cryo-EM density and are shown. The residue Tyr268 (Tyr254) has not been resolved in either structure. All structures were superimposed via *MatchMaker* in Chimera.

| | GSIII (EMD-26377, PDB 7U6O) |
|---|---|
| **Data collection** | FEI Titan Krios |
| Voltage (kV) | 300 |
| Spherical aberration (mm) | 2.7 |
| Total exposure dose ($e^-$ Å$^{-2}$) | 100 |
| Pixel size (Å) | 0.6509 |
| **Data processing** | |
| Software used | cryosparc v3 |
| Initial number of particles | 554,543 |
| Final number of particles | 270,133 |
| Symmetry imposed | C6 |
| Refinement | Homogeneous |
| Map resolution (Å) | 3.18 |
| FSC threshold | 0.143 |
| 3D reference model | *ab initio* |
| **Validation** | RealSpaceRefine |
| CC (volume) | 0.78 |
| Bonds (RMSD), length (Å) | 0.003 |
| Bonds (RMSD), angles (º) | 0.596 |
| Ramachandran favored (%) | 94.54 |
| Ramachandran allowed (%) | 5.46 |
| Ramachandran outliers (%) | 0.00 |
| Rotamer outliers (%) | 0.06 |
| Cß outliers (%) | 0.00 |
| Clashscore | 18.38 |
| MolProbity score | 2.14 |

Table S1. Cryo-EM data information and PHENIX refinement statistics for GSIII.
